# Microfabrication-based engineering of biomimetic dentin-like constructs to simulate dental aging

**DOI:** 10.1101/2023.09.10.557073

**Authors:** Simon Álvarez, Jose Morales, Paola Tiozzo-Lyon, Pablo Berrios, Valentina Barraza, Kevin Simpson, Andrea Ravasio, Xavier Monforte Vila, Andreas Teuschl-Woller, Christina MAP Schuh, Sebastian Aguayo

## Abstract

Human dentin is a highly organized dental tissue displaying a complex microarchitecture consisting of micrometer-sized tubules encased in a mineralized type-I collagen matrix. As such, it serves as an important substrate for the adhesion of microbial colonizers and oral biofilm formation in the context of dental caries disease, including root caries in the elderly. Despite this issue, there remains a current lack of effective biomimetic *in-vitro* dentin models that facilitate the study of oral microbial adhesion by considering the surface architecture at the micro- and nanoscales. Therefore, the aim of this study was to develop a novel *in-vitro* microfabricated biomimetic dentin surface that simulates the complex surface microarchitecture of exposed dentin. For this, a combination of soft lithography microfabrication and biomaterial science approaches were employed to construct a micropitted PDMS substrate functionalized with mineralized type-I collagen. These dentin analogues were subsequently glycated with methylglyoxal (MGO) to simulate dentin matrix aging *in-vitro* and analyzed utilizing an interdisciplinary array of techniques including atomic force microscopy (AFM), elemental analysis, and electron microscopy. AFM force-mapping demonstrated that the nanomechanical properties of the biomimetic constructs were within the expected biological parameters, and that mineralization was mostly predominated by hydroxyapatite deposition. Finally, dual-species biofilms of *Streptococcus mutans* and *Candida albicans* were grown and characterized on the biofunctionalized PDMS microchips, demonstrating biofilm specific morphologic characteristics and confirming the suitability of this model for the study of early biofilm formation under controlled conditions. Overall, we expect that this novel biomimetic dentin model could serve as an *in-vitro* platform to study oral biofilm formation or dentin-biomaterial bonding in the laboratory without the need for animal or human tooth samples in the future.

## 2. Introduction

Human teeth are highly specialized organs anatomically consisting of a crown and one or more roots (**Figure 1**). Furthermore, the tooth contains four distinct tissues including enamel, dentin, cementum (hard tissues), and pulp (soft and non-calcified tissue)^1^, amongst which dentin occupies the largest volume. Dentin is a bone-like mineralized tissue mostly comprised of hydroxyapatite (HA) crystals and an organic matrix dominated by type-I collagen^2,3^. Dentin shows an organized and complex microarchitecture consisting of micrometer-sized tubules (**Figure 1**) that can become exposed once dentin comes into contact with the environment including tooth wear, enamel erosion, or gum recession^4^. Once uncovered, the dentinal surface can act as a substrate of considerable clinical significance for the colonization by oral bacteria and fungi^5,6^.

**Figure 1:**
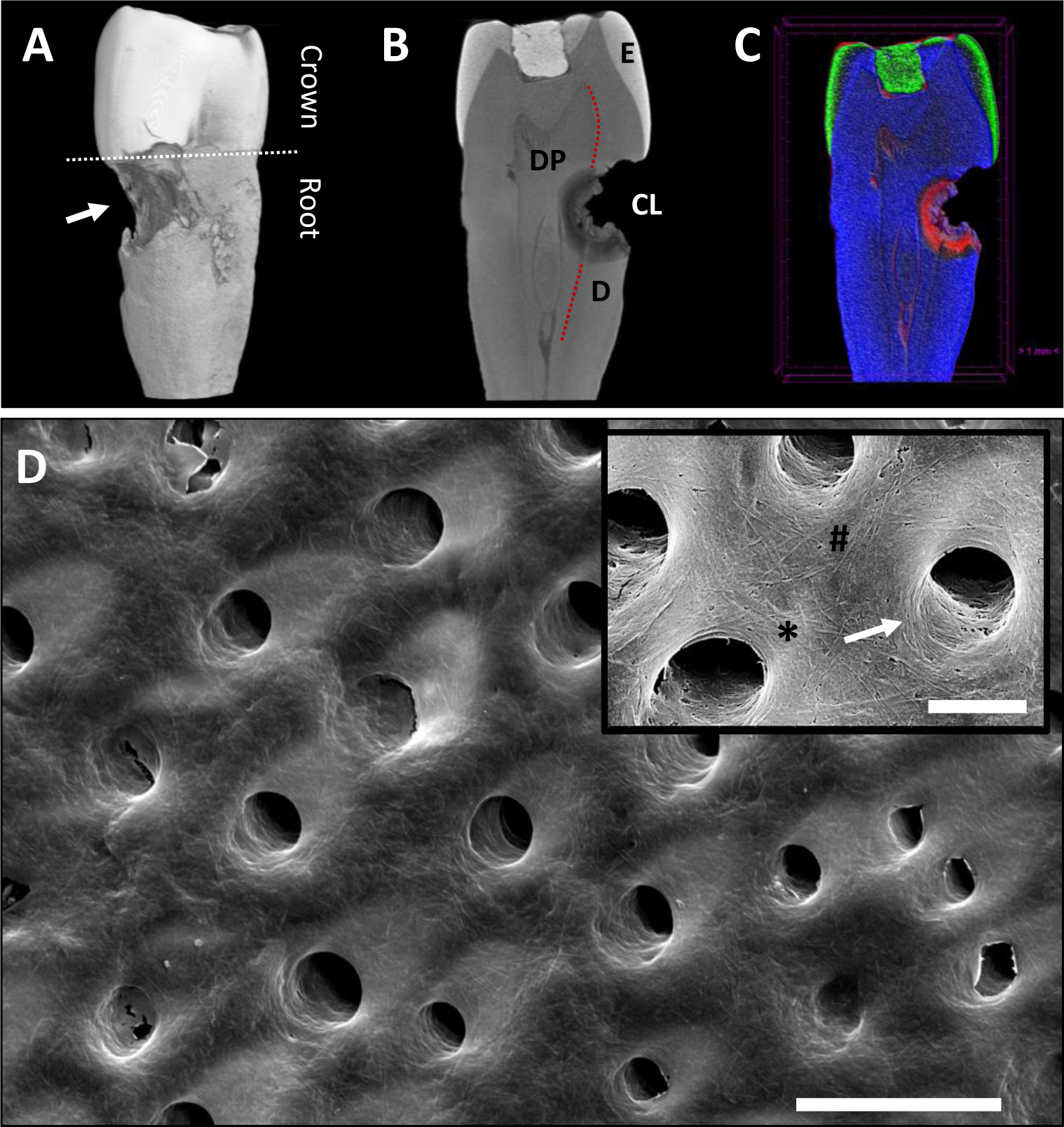
Dentin as a relevant mineralized tissue for biofilm formation and root caries. (A) Micro-CT 3D volume rendering of a tooth with an advanced root caries lesion (Arrow). (B) Micro-CT transversal slice through the middle section of the same tooth, where the enamel (EM), dentin (D), dental pulp (DP), and root caries lesion (CL) can be observed. (C) Micro-CT pixel intensity transform function (red: low intensity; blue: medium intensity; green: high intensity) that shows the important mineral loss associated with the caries lesion. (D) Scanning electron micrograph of demineralized dentin and dentinal tubules from a 62-year-old tooth (scale bar 10µm; inset 3µm). The heterogenicity of collagen morphology can be seen, with areas of increased deposition density (*), single individualized fibrils (#), and regions of circumferential collagen deposition around the tubules (arrow).

As a result, microbial adhesion to dentin is considered the initial step in the establishment of a complex biofilm that promotes root caries development – a highly prevalent chronic dental disease that affects the elderly^6,7^. As substrate characteristics (i.e., charges, roughness) are known to modulate biofilm formation, it is believed that changes in the biological and mechanical properties of dentin play an important role in bacteria-surface interactions. Within this context, it is currently known that during aging, important changes occur in the organic matrix of dentin such as the accumulation of advanced glycation end-products (AGEs) such as pentosidine and methylglyoxal (MGO)^3,8,9^. Glycation of tissues – including collagen - has been shown to modulate the adhesion of oral bacteria to tissues^10,11^ and therefore, understanding the early adhesion and biofilm formation on glycated dentin is instrumental in finding novel ways to prevent and treat root caries in the elderly.

Despite this clinical problem, effective biomimetic *in-vitro* approaches for studying oral microbial adhesion to dentin are lacking. Over the years, some models have been developed to simulate the intricate interactions occurring at the bacteria- dentin interface – including atomic force microscopy (AFM)-based studies - in an effort to comprehend the complexity of biofilm formation^10,12,13^. However, in many cases, the employed *in-vitro* models use amorphous substrates that despite considering some of the biological components, do not reproduce the micro- and nano-architecture of dentin. On the other hand, the use of *ex-vivo* dentin specimens from animals or human donors can be difficult to obtain due to practical and ethical limitations, as well as qualitative variations between different teeth and individuals^14,15^. Thus, efforts remain to generate effective *in-vitro* biomimetic constructs with microtopography and nanomechanical behaviors that are analogous to ex-vivo dentin specimens.

Nowadays, the use of microfabrication-derived approaches – such as soft-lithography - are slowly gaining traction in bioengineering as these techniques enable the development of accurately designed surfaces with micrometer and nanometer precision^16–19^. These techniques involve transferring a customized computer-generated micropattern onto a biomaterial substrate that serves as a template for subsequent modifications through processes such as photopolymerization or biofunctionalization^20^. One of the notable advantages of microfabrication is the capacity to precisely and reproducibly control three-dimensional micrometric geometries that can simulate complex in-vivo tissues in the laboratory^21^. Nonetheless, the use of microfabrication and biofunctionalization to construct dentin-like substrates for *in-vitro* use in biofilm formation assays has not been explored to date.

Therefore, this study aims to develop a novel *in-vitro* microfabricated biomimetic dentin surface to simulate the complex surface microarchitecture of exposed dentin, to be used as a model for the study of bacterial adhesion and biofilm formation in the context of dentinal aging. We hypothesize that these biomimetic dentinal constructs will display similar characteristics to native dentin and serve as an effective *in-vitro* model to characterize clinically relevant biofilm formation in the context of tooth aging.

## 3. Methods

### 3.1. Tooth collection and root caries imaging

Teeth for this study were obtained under informed consent and local ethical approval (#210615001). Following extraction, teeth were debrided from soft tissues and decontaminated in 70% ethanol for 72 h, washed with ultrapure water (dH_2_O), and imaged with a SkyScan 1272 Micro Computed Tomography (micro-CT) system (Bruker, USA). Micro-CT scans were processed in the CT Vox software v.3.3.1 (Bruker, USA) in order to obtain 3D renderings and digital slices of the tooth structure (**Figure 1**). For scanning electron microscopy (SEM), images of dentin ultrastructure, 100µm thin tooth sections were prepared with an SP1600 hard tissue microtome (Leica Biosystems, USA), and were submerged in a 37% phosphoric acid solution (Sigma-Aldrich, USA) for 30s, followed by a 10s rinse in sodium hypochlorite (Sigma-Aldrich, USA), washed three times with dH_2_O and air-dried. Subsequently, the sections were submerged in a solution of 2.5% glutaraldehyde for 24 h, dehydrated with an increasing 25%, 50%, 70%, 90%, 100% ethanol series, gold sputter-coated and imaged with a Field Emission Scanning Electron Microscope (FESEM, Quanta FEG250, FEI, USA).

### 3.2. Substrate design and photolithography

Firstly, a pattern consisting of 5 µm diameter circles with separation distances of 8 µm was created in the KLayout software (Matthias Köfferlein, version 0.27.12) and used to create SU-8 micropatterned silicon wafers through photolithography. Briefly, a negative SU-8 photoresist (GM1040, Gersteltec) was added to a silicon wafer and then placed in a spin coater to generate a thin SU-8 film of approximately 5 µm. Subsequently, the computer-designed micropattern was directly written onto the photoresist by exposure to UV light with a laser pattern generator (Heidelberg Instruments MLA 100). As the photoresist polymerizes only in the areas exposed to UV light, the unexposed portions were removed using a SU-8 developer, leaving only the patterned pillar design adhered to the wafer (**Figure 2**).

**Figure 2:**
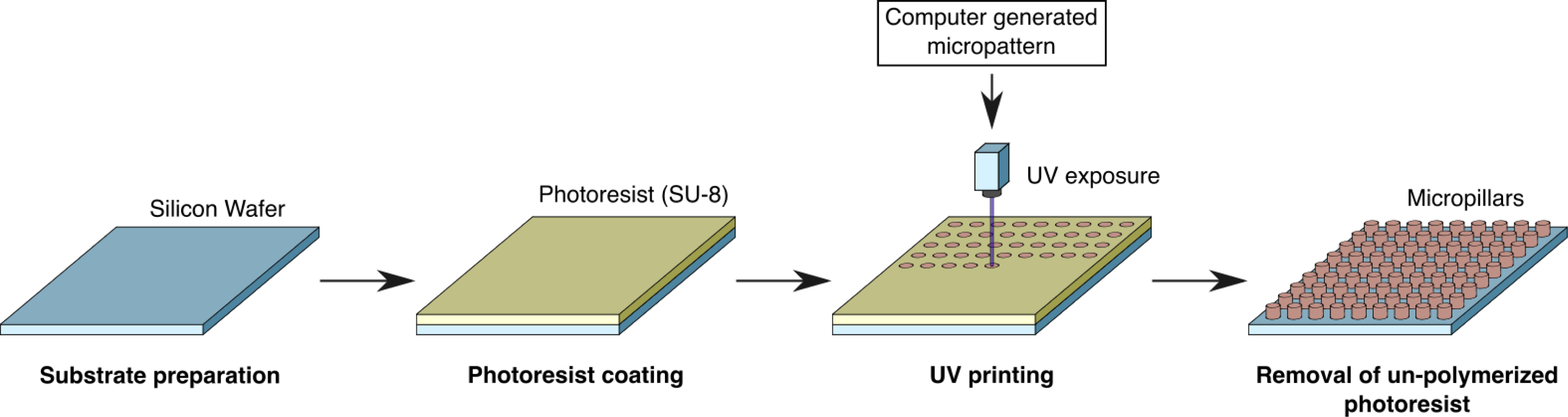
Schematic representation of the photolithography microfabrication pipeline.

### 3.3. PDMS substrate microfabrication

Following the fabrication of the master mold, PDMS (Poly-dimethyl-siloxane; SYLGARD® 184, Dow-Chemical) was prepared following the specifications provided by the manufacturer. Briefly, the PDMS base and crosslinker were mixed in a proportion of 10:1, respectively and carefully deposited over the micropatterned silicon wafers, ensuring uniform coverage of the surface. Subsequently, the PDMS-coated wafer was incubated at 80°C for 1 h and subsequently cooled to room temperature. Using a disposable biopsy punch, the PDMS mold was cut into 4 mm diameter specimens and air plasma-cleaned for 60 s to activate functional groups on the PDMS surface. The surfaces were then coated with APTES by vapor deposition using approximately 100 µl of hydrolyzed APTES 97% ((3-aminopropyl)triethoxysilane; Sigma-Aldrich) in a vacuum chamber for 2 h.

### 3.4. Microfabricated wafer and PDMS substrate characterization

Following PDMS microfabrication, substrates were characterized with a Panthera U light microscope (Motic, China) at 4x and 40x magnification to confirm pattern structure and regularity. Furthermore, micropatterns were imaged with a Field Emission Scanning Electron Microscope (FE-SEM, Quanta FEG250, FEI, USA) at varying inclination angles with 1500x, 2500x and 10000x magnification. Finally, both the micropillar diameter and separation were quantified (**Figure 3D**) from FESEM images by using the FIJI open-source software (ImageJ).

**Figure 3:**
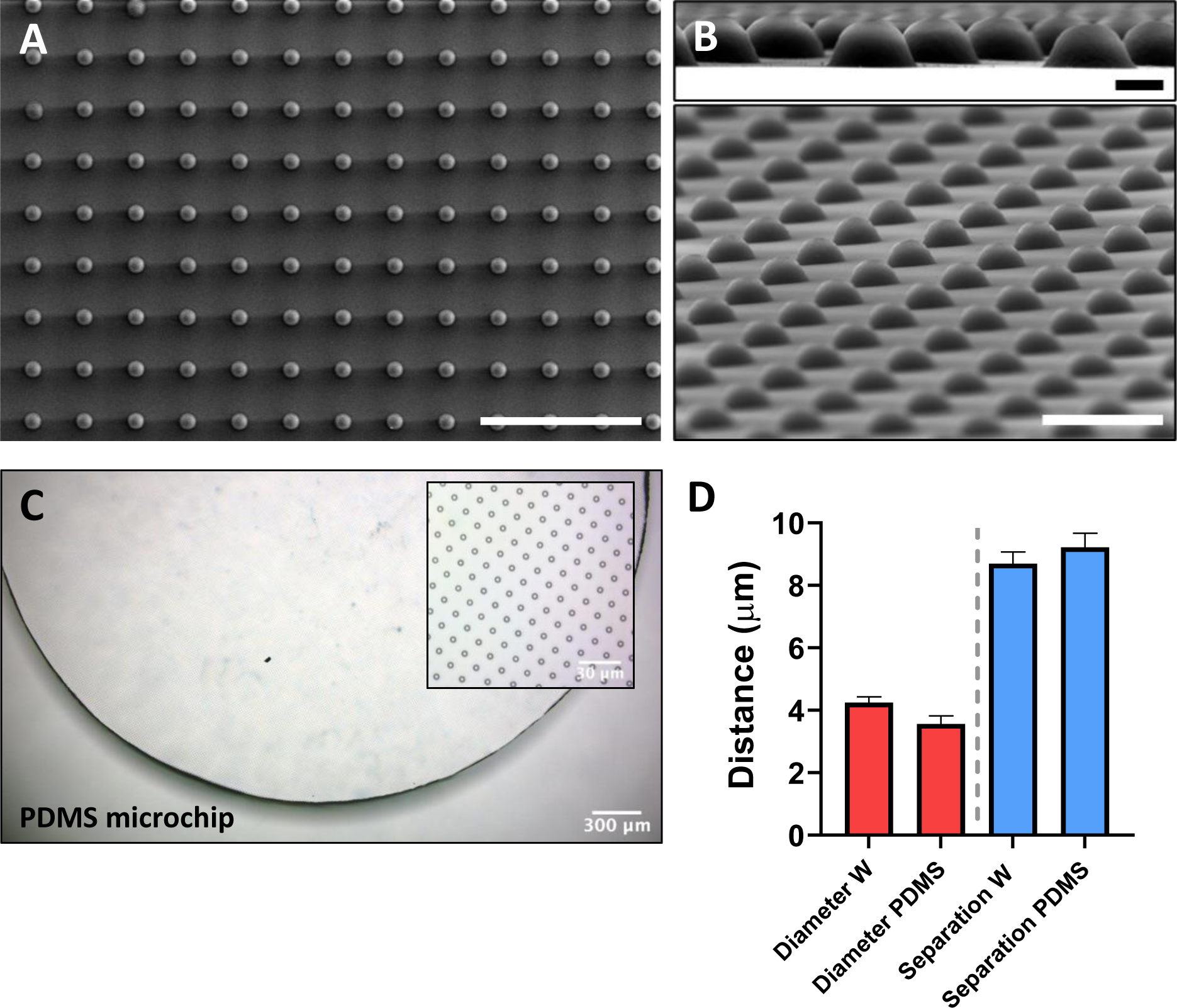
Microfabricated silicon wafers and PDMS surfaces with tubule-like morphology. (A) Scanning electron micrograph of SU-8 micropillars after photolithography microfabrication (Scale bar: 40µm). (B) Lateral views of the SU-8 micropillar array (Top scale bar: 2µm; Bottom scale bar: 10 µm). (C) Optical microscopy of the resulting PDMS chip displaying the tubule-like morphology of the substrate in a regular array pattern. (D) Micropit diameter and separation lengths for both the wafer (W) and PDMS constructs (n=300 for micropits, n=295 for separations; across three substrates each).

### 3.5. PDMS substrate functionalization with type-I collagen

To prepare the collagen solution, 330 µl of rat tail type-I collagen (3 mg/ml; Gibco A1048301) was gently diluted in 660 µl 1X PBS for a final collagen concentration of 1 mg/ml. The prepared collagen solution was deposited onto the PDMS mold, covering the whole surface, and incubated at 37°C for 1 h to allow fibril formation. Collagen-coated PDMS was imaged with phase contrast microscopy and FESEM as previously described above, and collagen deposition was confirmed with Attenuated Total Reflectance-Fourier Transform Infrared Spectroscopy (ATR-FTIR) by employing an IRTracer 100 (Shimadzu, Japan).

### 3.6. Mineralization of collagen coated PDMS surfaces

To prepare the mineralization solution, the protocol described by Tas and Bhaduri was followed^22^. Briefly, a stock solution of NaCl, KCl, CaCl_2_ ·2H_2_O, MgCl_2_ · 6H_2_O, and NaH_2_PO_4_ was prepared in ultrapure water. To activate the mineralization process, 84 mg of NaHCO_3_ was added to 100 ml of solution at room temperature during magnetic stirring. Subsequently, the PDMS specimens were placed into a 96-well plate and coated with 100µl of the mineralizing solution at room temperature for 30 min. Finally, PDMS substrates were washed twice with 1X PBS. Calcium deposition was confirmed with Alizarin Red staining, both with phase contrast microscopy and quantification using a micromodal plate reader at 450nm (Synergy HT, Biotek) (**Supplementary** Figure 1).

### 3.7. Analysis of mineralization with X-ray diffraction (XRD)

To determine the crystallographic structure of substrate mineralization, XRD was performed on the mineralized collagen-PDMS samples. For XRD, the resulting mineralizing precipitate formed on substrates was washed three times with 50ml of 1X PBS, filtered using filter paper, and dried at RT. Samples were analyzed with a D8 Advance diffractometer (Bruker, USA) to determine the crystallographic structure of the resulting mineralization.

### 3.8. *In-vitro* aging of microfabricated mineralized collagen substrates

To replicate collagen aging, the resulting microfabricated collagen surfaces were incubated with 10 mM methylglyoxal (MGO, Sigma Aldrich) for 48 h at 37°C. Subsequently, samples were carefully washed three times with 1X PBS to remove any residual MGO from the substrate and kept in 1X PBS until experimentation. To quantify the modification of mineralized collagen by MGO, 1 mg/ml collagen gels were incubated with 10mM MGO and the clinically relevant MGO-derived advanced glycation end-products (AGE) hydroimidazolone MG-H1 was measured using a competitive ELISA assay (Abcam, UK) according to the manufacturer’s instructions and previously published approaches^23^.

### 3.9. Atomic force microscopy (AFM) imaging and nanomechanical force-mapping of mineralized substrates

Topographic and force-curve measurements were performed in air with a MFP 3D- SA AFM (Asylum Research, USA) utilizing SCOUT 350 RAu cantilevers with a nominal spring constant of 42 N/m (NuNano, UK). Individual tip calibration was performed as described by the manufacturer using a clean glass cover, and cantilever tuning was performed at a target amplitude of 1V. Topographic images were made for all materials in AC mode at a scan size of 20x20 µm, a resolution of 256 points and lines, and a scan rate of 0.8 Hz. The set point and integral gain was adjusted for each sample to achieve higher resolution, ranging from 200 mV for bare PDMS and 500 mV for functionalized PDMS. The force-mapping measurements were made in a scan size of 20x20µm with 16X16 and 32X32 points for PDMS and mineralized PDMS surfaces, respectively. A force distance of 5 µm and a velocity of 9.92µm/s was used for all force-curve measurements.

Young’s modulus was obtained using the Johnson, Kendall, and Robert model in the Asylum Research proprietary software. Three scans were obtained in representative areas for each condition. All data and images were obtained and analyzed using the Asylum Research proprietary software (v.16.10.211) and histograms were generated in the IGOR 8.04 software.

### 3.10. Analysis of mineralized microfabricated substrate ultrastructure with FESEM and EDX

Ultrastructural imaging of microfabricated PDMS substrates was carried out with a Quanta FEG250 FESEM. Samples were fixed with PFA 4% at RT for 30 min, dehydrated in a series of ethanol solutions of increasing concentration (25, 50, 70, 90, and 100% ethanol), followed by hexamethyldisilazane (Sigma) drying for 5 min, and subsequent airdrying. Finally, samples were sputter-coated with 10nm gold and visualized with 10kV acceleration voltage, and surface composition was explored with Energy-dispersive X-ray spectroscopy (EDX) using the FESEM- coupled Octane Pro Silicon Drift Detector system (EDAX, USA).

### 3.11. Microbial cultures and dual-species biofilm formation

*C. albicans* (90028) and *S. mutans* UA 159 strains were maintained and incubated on solid TSB (Tryptic Soy Broth) or BHI (Brain Heart Infusion; BD Bioscience, San Jose, CA, USA) agar plates for 24 h at 37°C and 5% CO_2_. Cell numbers for *C. albicans* and *S. mutans* were standardized using calibration curves to a microbial concentration of 5 x 10^6^ cells/ml. *C. albicans* and *S. mutans* were seeded onto the mineralized collagen substrates in a proportion of 1:1 bacteria (12.5x10^2^ cells of *C. albicans* – 12.5x10^2^ cells of *S. mutans*) and grown in a media containing BHI and TSB (1:1) at 37°C and 5% CO_2_ for 24 h.

### 3.12. Analysis of dual-species biofilm formation by epifluorescence and confocal microscopy

The resulting dual-species biofilms were characterized with epifluorescence and confocal microscopy. After a 24-hour incubation, biofilms were gently washed with 1X PBS to remove unattached cells and stained with a solution of SYTO9 (Live/Dead Baclight, ThermoFisher, USA), and CalcoFluor White (Sigma Aldrich, USA) in BHI for 30 min at RT in darkness. Finally, samples were washed twice with 1X PBS and PDMS discs were mounted onto glass coverslips for microscopy analysis utilizing a Zeiss LSM 880 confocal microscope with Airyscan detection, utilizing the DAPI (Ex/Em: 365/445) and GFP (Ex/Em: 470/525) channels for CalcoFluor White and SYTO9, respectively, at 20x or 40x. For epifluorescence microscopy, images were obtained with a Nikon Ti2 epifluorescence microscope equipped with a Hamamatsu high-resolution camera at 20x and 40x **(Supplementary** Figure 2).

### 3.13. Statistical analysis

All quantitative data was analyzed with GraphPad Prism v9 and Asylum Research Igor software. Data is presented as means ± standard deviation, or as a histogram of resulting values. After determining parametric or non-parametric distribution either t-test, Mann-Whitney, or one-way ANOVA with Tukey’s multiple comparisons test with were performed considering significance at p<0.05.

## 4. Results and discussion

### 4.1. Construction of a micropitted collagen-coated PDMS substrate to simulate the dentinal microarchitecture

As previously described, the complex microarchitecture of human dentin involves a mineralized collagen matrix organized in a microtubular fashion, with areas of amorphous fibrillar anatomy as well as circumferential collagen fibril arrangement (**Figure 1D**). Once dentin is exposed to the oral environment, it becomes a relevant surface for the attachment of oral microbes and biofilm, as well as a relevant surface for the adhesion of restorative dental biomaterials during clinical treatment. Therefore, to replicate this superficial morphology in-vitro, we have utilized microfabrication to generate a micropillar array from which a PDMS negative could be cast via soft-lithography (**Figure 3A-B**). This resulted in the construction of a PDMS surface with micropit diameters of ∼3.5µm and separations of ∼8.5µm (**Figure 3C-D**), similar to the dimensions found in some regions of native dentinal tissue^2^. For ease of construction, biopsy punches were utilized to generate 4mm diameter PDMS discs that can be placed into 96-well plates to allow for high-throughput processing (**Figure 3C**). Furthermore, the surface morphology and elastic properties at the micro- and nanoscale were explored with AFM imaging and quantitative force-mapping (**Figure 4**). These results confirmed the micropit dimensions and arrays observed with light and electron microscopy, and the Young’s modulus of the PDMS construct was found to be in the low MPa range, consistent with previous reports in the literature for similar substrates^24–26^.

**Figure 4:**
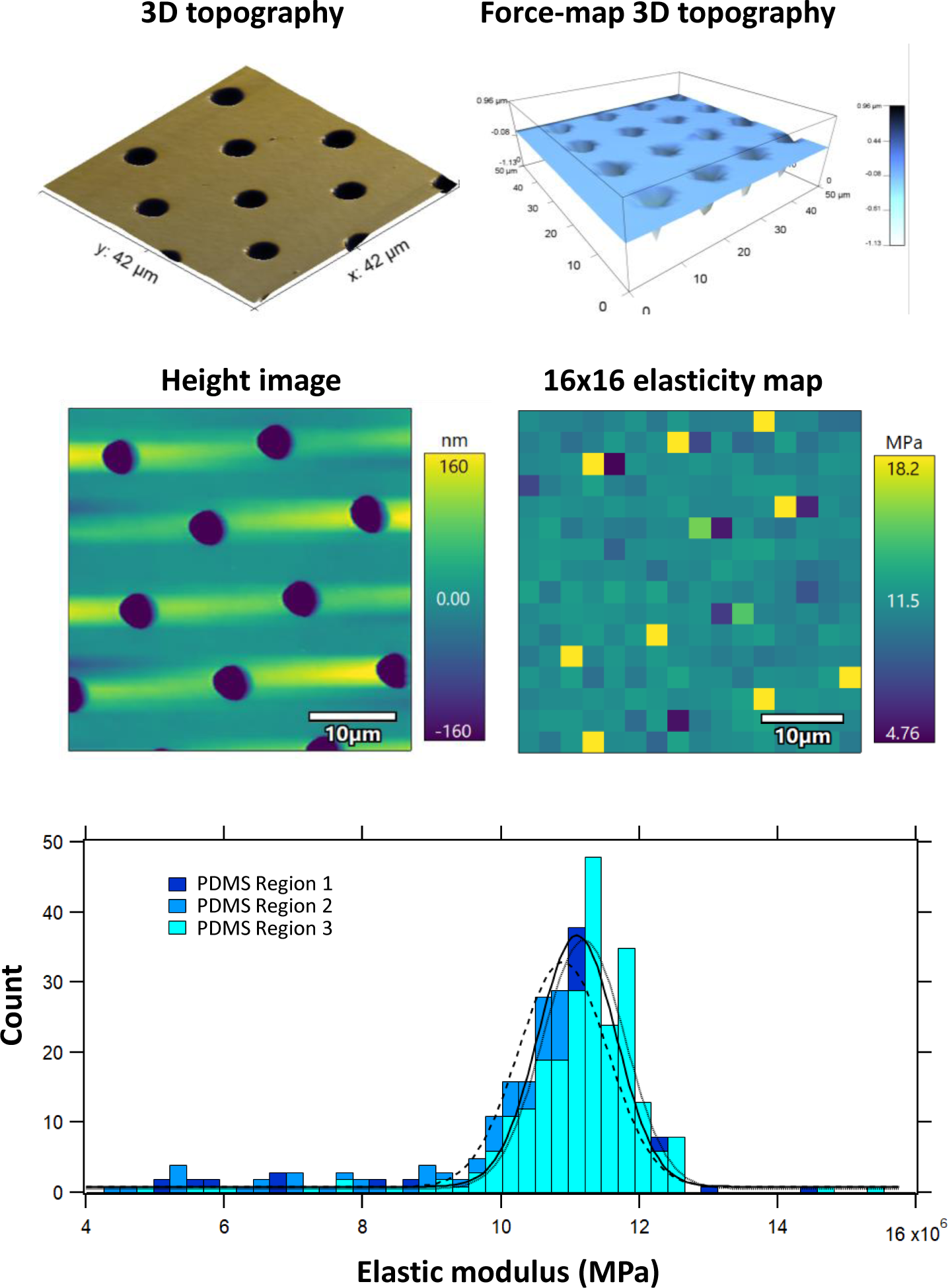
Nanomechanical and topographical characterization of PDMS substrates with atomic force microscopy (AFM). AFM topography (upper left) and 3D force-map topographical rendering (upper right) of the PDMS substrates, demonstrating the micro- and nanoscale architecture of the micropit array. 16x16pixel force-maps were utilized to obtained nanomechanical elastic properties from the substrates, which displayed a reproducible elasticity in the low MPa range.

Once the microfabricated PDMS substrates were created, the next step was to modify the surface with type-I collagen in order to emulate the main component of the dentinal collagen matrix. This was obtained by pretreating the PDMS surface with plasma cleaning and subsequently surface coating with APTES. After incubation, a thin layer of amorphous type-I collagen fibrils could be observed coating the surface of the PDMS construct with FESEM and phase contrast microscopy (**Figure 5A-C**). The mechanical tensions generated by the collagen matrix result in micropit diameter heterogeneities that are reminiscent of the anisotropic behavior of in-vivo biological tissues (**Figure 5A**). High magnification SEM imaging demonstrated that collagen also organized circumferentially around the micropits, as well as at different fibril densities across the inter-pit regions (**Figure 5B**). This arrangement is very similar to the native organization of the dentinal collagen matrix observed *in-vivo* (**Figure 1D**)^3^. Furthermore, ATR-FTIR confirmed the presence of a collagen pellicle on the surface following functionalization, confirmed by the presence of the characteristic amide bands as previously observed in literature (**Figure 5D**) ^23,27^.

**Figure 5:**
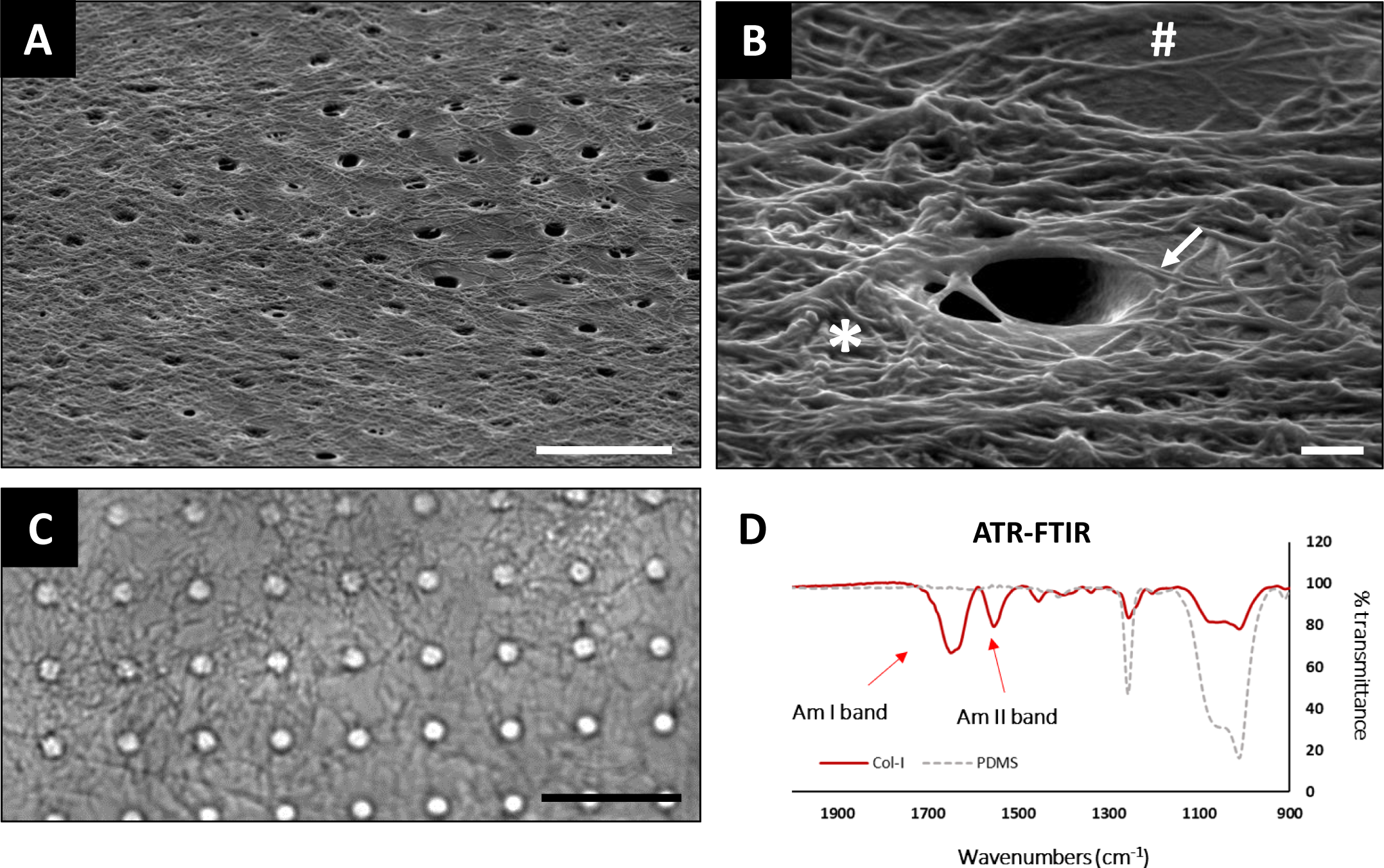
Biofunctionalization of PDMS micropitted substrates with fibrillar type-I collagen. (A) Scanning electron micrograph of a PDMS-collagen construct showing the overall surface morphology (Scale bar: 20µm). (B) High-magnification scanning electron micrograph of the type-I collagen layer showing areas of increased deposition density (*), individualized fibril regions (#), and circumferential collagen deposition around the micropits (arrow), similar to the in-vivo heterogenicity of dentin (Scale bar: 1µm). (C) Phase contrast microscopy in PBS demonstrates the fibrillar organization of type-I collagen on the PDMS micropit array (Scale bar: 20µm). (D) ATR-FTIR spectra for bare PDMS and collagen- coated PDMS. Note the increase in Am-I and Am-II band intensities confirming the deposition of collagen on the surface.

### 4.2. Mineralization and development of a biomimetic in-vitro dentinal aging model

Subsequently, the collagen-coated PDMS constructs were immersed in a previously described mineralizing solution^22^ in order to generate HA deposition within the collagen matrix (**Figure 6A**). EDX characterization of mineralized PDMS discs demonstrated surface deposition of Ca, P, and O; and XRD showed that the main mineral corresponded to HA followed by chlorapatite (**Figure 6B and 6C**).

**Figure 6:**
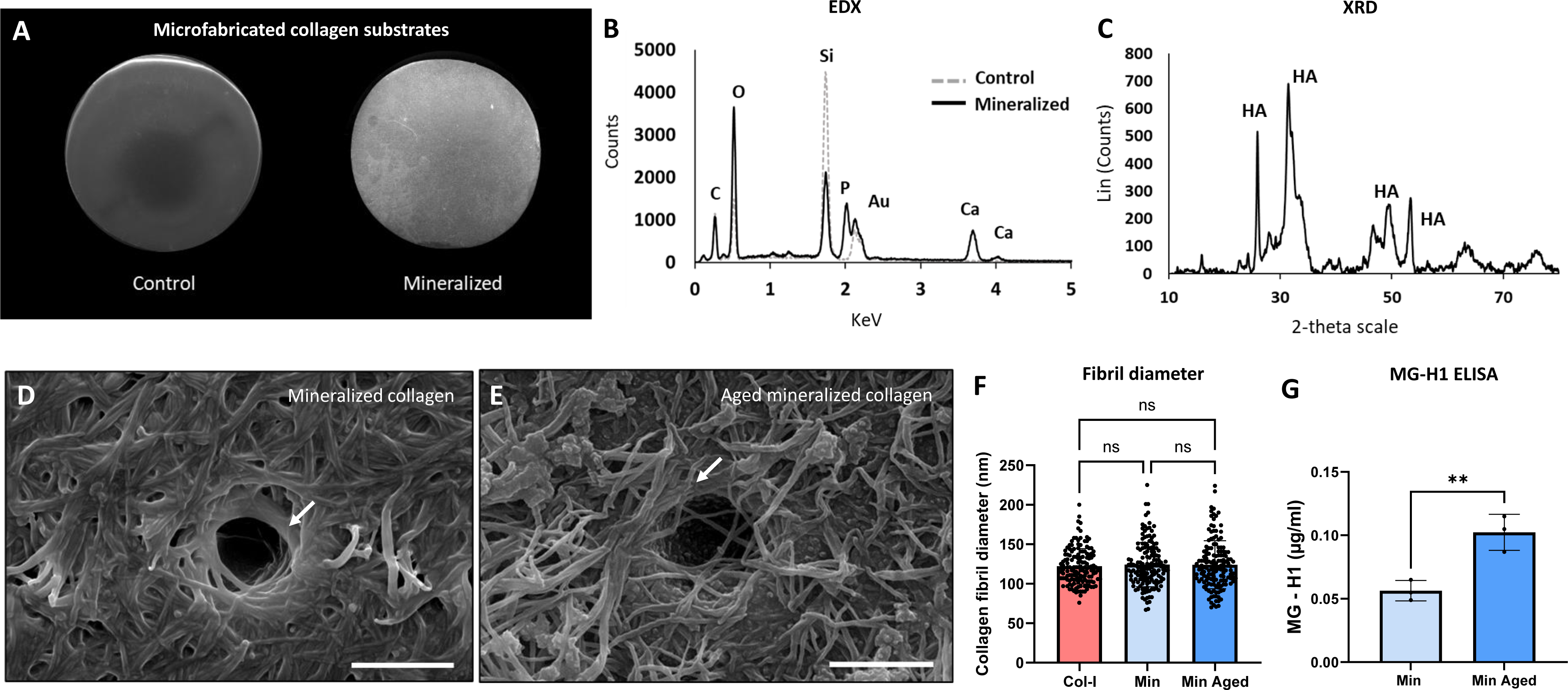
Mineralization and *in-vitro* aging of biomimetic dentin constructs. (A) Scanning electron micrographs of non-mineralized and mineralized 4mm diameter collagen-PDMS constructs. (B) energy-dispersive X-ray and (C) X-ray diffraction analysis of PDMS constructs confirming HA-dominant mineralization. (D) and (E) Scanning electron micrographs of mineralized constructs before and after aging with methylglyoxal (MGO), respectively (Scale bars: 3µm). (F) Collagen fibril diameters for non-mineralized, mineralized, and aged mineralized substrates displayed no statistical differences among conditions (n=150 collagen fibrils per condition; ns: non-significant, one-way ANOVA withTukey’s multiple comparisons test). (G) ELISA detection of the MGO-derived advanced glycation end-product MG-H1, demonstrating a significant increase in glycation following *in-vitro* aging of mineralized collagen-PDMS constructs (n=3; **p<0.01, t-test).

SEM imaging demonstrated that the collagen network did not lose its morphology and that the short mineralization time used in this study was not able to completely encase the collagen network (**Figure 6D**). Comparable results were found by other research groups utilizing similar mineralizing solutions at short incubation times, confirming the utility of this approach for obtaining biologically relevant mineralized tissue analogs *in-vitro* ^28,29^. In this particular work, the ability to generate microfabricated, mineralized collagen-coated surfaces is of instrumental importance for the development of *in-vitro* biomimetic dentin substrates, and serves as a novel platform to create biologically-relevant dentin analogs for *in-vitro* experimentation.

It is currently known that *in-vivo*, dentin goes through age-related alterations such as the accumulation of AGEs within the organic matrix such as pentosidine and MG-H1^3,8,9^. Therefore, it is of interest to determine if AGEs can also be incorporated into microfabricated mineralized collagen matrices *in-vitro*. For this, the mineralized chips were incubated in 10 mM MGO as previously described^10^ for 48 h in order to allow glycation of collagen. SEM imaging showed that MGO did not alter the organization of the collagen scaffold and that the micropit dimensions were maintained following incubation (**Figure 6E**). No significant differences were found regarding the diameter of collagen fibrils between the non-mineralized, mineralized, and aged-mineralized groups, with most fibrils displaying diameters between 100-150 nm (**Figure 6F**). Most importantly, the glycation of mineralized collagen with MGO was confirmed through the incorporation of the MGO-derived AGE MG-H1 into the substrate (**Figure 6G**). This AGE has been found in dentinal and periodontal samples of older individuals and is known to be relevant and associated with tissue aging and diseases such as diabetes mellitus^3,30–32^.

### 4.3. Nanomechanical properties of the novel dentin biomimetic constructs probed by AFM

As a next step, the nanomechanical properties of the microfabricated *in-vitro* dentin constructs were explored with AFM-based force mapping (**Figure 7**). Overall, the mineralized collagen surfaces were found to have an elastic modulus below ∼0.5 GPa; however, the Young’s modulus significantly increased after glycation with MGO. This is similar to what has been reported in previous literature, where glycation with AGEs has been seen to alter the mechanical properties and increase the Young’s modulus of *in-vitro* collagen constructs, dentin, and bone^3,23,33,34^. Also, some heterogenicity was observed in the elastic properties of the mineralized collagen substrates across the surfaces, probably due to the presence of regions with different amounts of collagen deposition as well as heterogeneous mineralization. Nevertheless, the elastic modulus of mineralized collagen both with and without MGO incubation was found to be over one order of magnitude higher than the bare PDMS microfabricated surface (**Figure 4**), confirming that the nanomechanical properties of the final biomimetic constructs are provided by the mineralized collagen layer on its surface.

**Figure 7:**
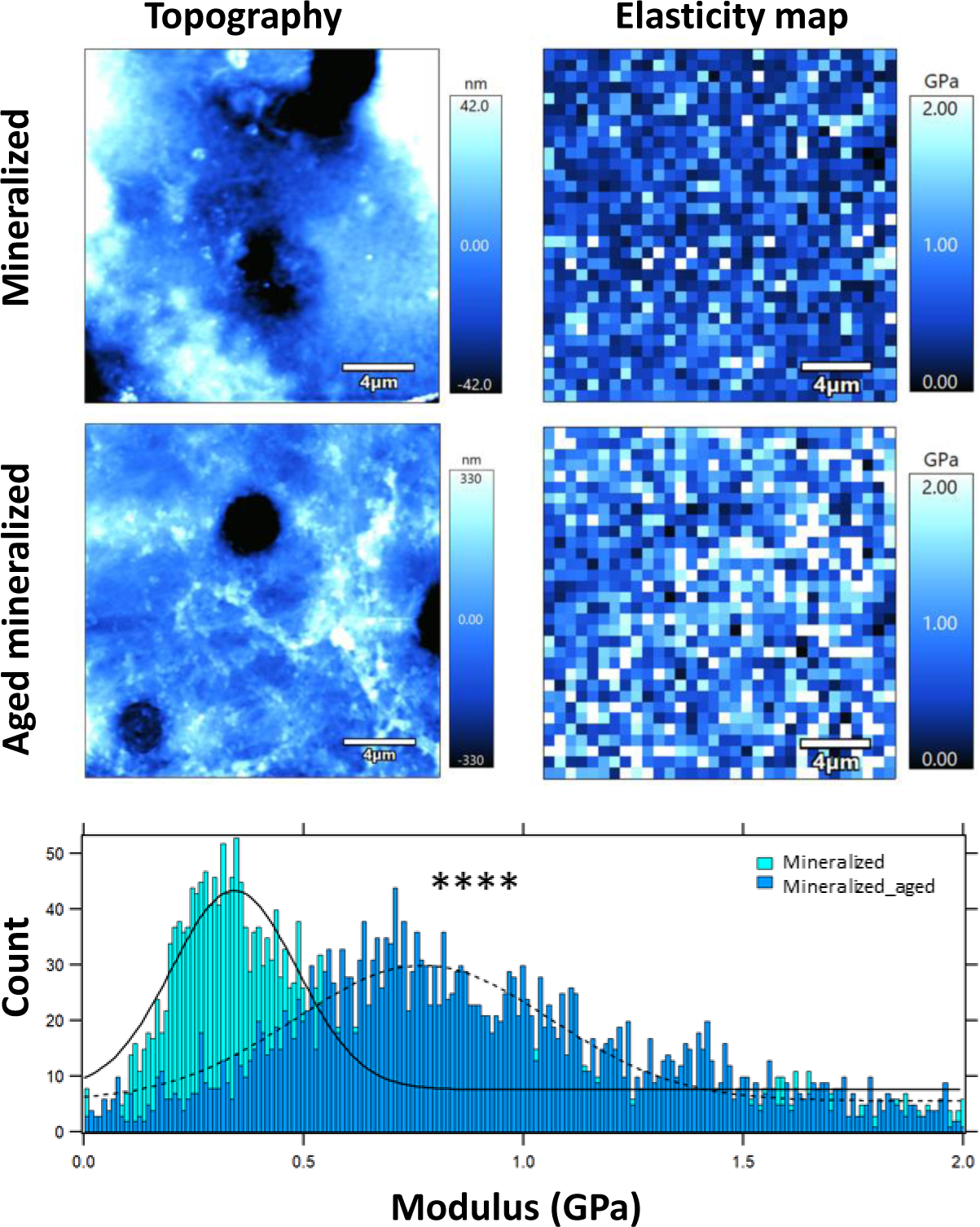
Nanomechanical characterization of biomimetic dentin constructs with AFM imaging and force-mapping. Topographical height images (left) and elasticity maps (right) for non-aged and aged mineralized constructs. Histogram showing the dispersion of Young’s modulus across the studied samples (n=3072 force curves; t-test).

### 4.4. Dual-species oral biofilm formation on microfabricated mineralized surfaces

During the process of dental caries – and particularly root caries – oral microbes can attach directly to dentin and are associated with the development and progression of biofilms^35,36^. During this process, species such as *S. mutans* and *C. albicans* play an important role in the pathogenesis of caries through a range of virulence factors such as collagen-binding activity, acidification of the environment, and extracellular polysaccharide secretion^37,38^. Therefore, there is a pressing need to study the early-stage adhesion of these species onto dentin surfaces – especially aged dentin - to develop novel preventive and treatment approaches against dental caries.

Within this context, the formation of a dual-species *S. mutans* and *C. albicans* biofilm on the surface of mineralized collagen-coated PDMS was explored with fluorescence microscopy. The PDMS disc format facilitated the manipulation of the substrates and allowed for the system to be incubated, marked, and placed onto the microscopy stage without ever being removed from buffer conditions. With this approach, the surface morphology of living biofilms can be characterized with phase contrast and fluorescence microscopy (**Figure 8**). Following a 24-hour incubation period, the formation of an intricate dual-species biofilm was observed, showing the presence of *C. albicans* (including hyphae) interacting with *S. mutans* on the surface of the biomimetic devices^39^. High-resolution transmitted images and fluorescence orthogonal z-stack reconstructions for Calcofluor White and SYTO9 were obtained, which allowed for the characterization of biofilm biomass morphology with high resolution (**Figure 8A**). Further z-stack 3D rendering of high-magnification dual-species biofilm was also acquired (**Figure 8B**), confirming that the microfabricated constructs are viable for the study of microbial-surface interactions in the context of early-biofilm formation. Finally, the formation of *S. mutans* and *C. albicans* biofilms on non-glycated and glycated substrates was confirmed, demonstrating the potential of using this platform to study the interaction between oral colonizers and dentin-like substrates in vitro (**Figure 8C**).

**Figure 8:**
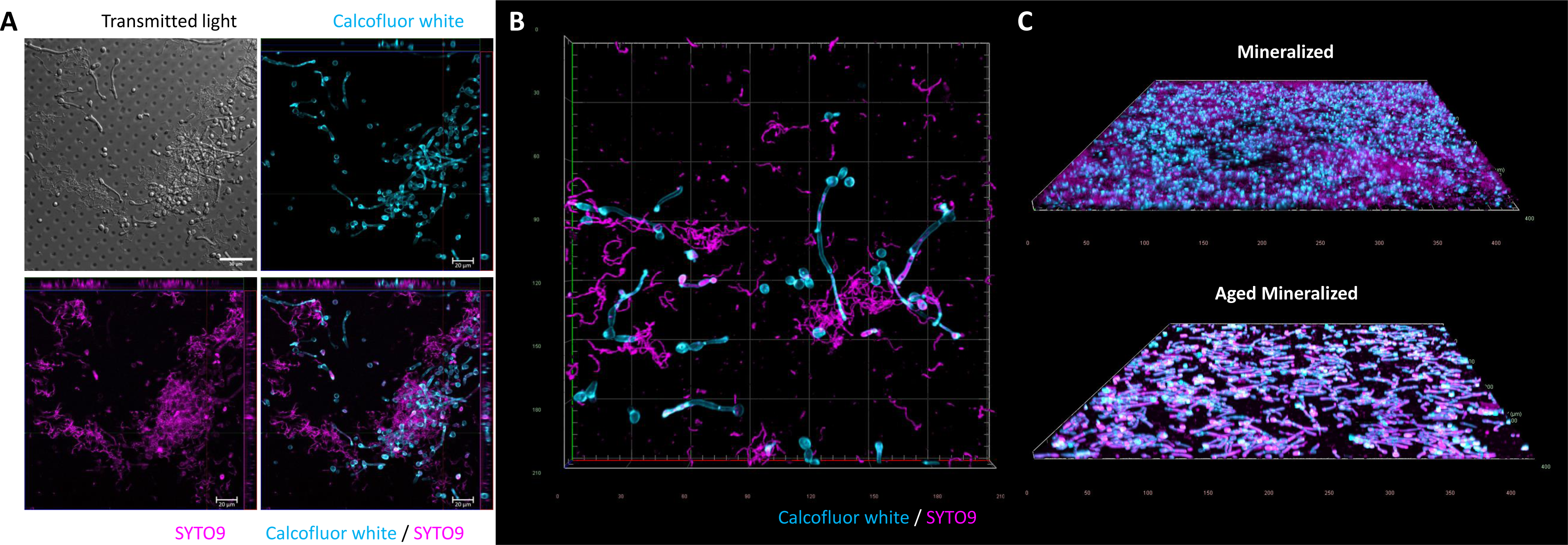
Proof-of-concept for dual-species biofilm formation on biomimetic dentin surfaces. (A) High-resolution transmitted light, Calcofluor White, SYTO9, and merged channels for a dual *Candida albicans* and *Streptococcus mutans* biofilm on a mineralized PDMS substrate. (B) High-resolution 3D rendering of a dual-species biofilm showing the interaction between fungal and bacterial cells on top of the microfabricated substrate. (C) 3D renders for dual-species biofilms formed on mineralized and aged-mineralized biomimetic substrates, displaying surface-specific morphological differences.

### 4.5. Toward an in-vitro biomimetic dentin microfabricated substrate

Currently, the use of microfabrication-derived approaches is slowly gaining traction in the field of dentistry, due to their ability to create environments that mimic cell, tissue, and organ-specific contexts^16–18,40–42^. Nevertheless, the ability to create *in-vitro* biomimetic constructs simulating the complex architecture of dentin has been difficult to accomplish to date. In this work, we have designed and microfabricated a novel dentin-like analog consisting of a mineralized type-I collagen matrix arranged in a micropit array simulating the tissue microarchitecture for the study of early oral biofilm formation. The nanomechanical properties of the organic matrix were obtained via the use of PDMS substrates functionalized with collagen fibrils, and the further mineralization with an HA-predominant phase has resulted in a final substrate with an elastic modulus in the high MPa/low GPa range characteristic for the native organic matrix of teeth^3,27,43,44^. Furthermore, the design of these surfaces in 4 mm diameter PDMS discs allows for its placement inside 96-well plates to carry out high-throughput experiments ranging from oral biofilm formation to potential tooth-biomaterial adhesive characterization, amongst others. Also, we have shown that it is possible to further tailor the mechanical and biological properties of these constructs (i.e., via glycation with relevant biological molecules) to study relevant processes *in-vitro* such as tooth aging and/or the effect of chronic hyperglycemia on dentinal properties^45,46^. Despite many advances in recent years, many questions remain regarding the pathogenesis of dental caries, particularly associated with vulnerable groups such as the elderly. Therefore, current dental research should focus on understanding how age-associated changes such as AGE accumulation within dental and periodontal tissues can promote disease^47^. Using this newly proposed model, we have preliminarily observed morphological differences in early-biofilm formation of a dual *S. mutans-C. albicans* biofilm on glycated surfaces, although a more in-depth characterization of these changes is expected to be carried out in future work.

Likewise, the development of organ-on-a-chip and tooth-on-a-chip technologies are opening new avenues towards the use of microfluidics for the study of important oral and dental processes^12,40,48^. Within this context, our novel in-vitro dentin constructs could easily be paired to microfluidic chambers and setups in order to e.g., explore single-bacterial adhesion of relevant oral microbial colonizers in real time with high spatiotemporal resolution and varying environmental factors (i.e., pH, sucrose availability, flow, etc.)^49^. The ability of this system to be paired with epifluorescence or confocal microscopy setups (**Figure 7 and Figure 8**) further strengthen its feasibility for the characterization of a diverse number of biological events involving the dentinal matrix and its interaction with biomaterials, cells, or microbes. As dental caries continues to be one of the most prevalent oral diseases and a significant contributor to tooth loss in populations worldwide^50^, the use of biomimetic microfabricated models such as this could pave the way towards novel ways to understand and treat this disease in the future.

## 5. Conclusion

The use of microfabrication, soft-lithography, and biofunctionalization proved to be effective tools to construct *in-vitro* biomimetic dentin surfaces consisting of a mineralized fibrillar type-I collagen matrix with a micropit surface array. Furthermore, these substrates were susceptible to biologically-observed ECM aging via MGO-derived glycation in order to model the process of AGE formation in teeth in the laboratory. These constructs were found to have topographical and nanomechanical properties similar to native dentin, with elasticity values in the 0.3- 2 GPa range. Finally, these biomimetic substrates were able to act as surfaces for the adhesion and growth of relevant dual-species oral biofilms of *S. mutans* and *C. albicans*, with results showing different three-dimensional microarchitectures for biofilms formed on both non-aged and aged substrates. Overall, this novel *in-vitro* biomimetic dentinal model could serve as a system to study oral biofilm formation or dentin-biomaterial bonding in the laboratory without the need for animal or human tooth samples in the future.

## Acknowledgements

This work has been supported by ANID FONDECYT #1220804, #1210872, and #1220803, ANID/SCIA/ACT192015, Millennium Science Initiative Program NCN19_170D, and ANID FONDEQUIP EQM140055, EQM180009, and EQM210101. Furthermore, we would like to thank the members of the City of Vienna Competence team “AgingTissue” (MA23#29-07) for the fruitful discussion on ECM changes during aging. Finally, authors would also like to thank the Advanced Microscopy Unit (UMA UC) for their support with the confocal imaging of dual-species biofilms.

## 6. Author contributions

SiA, JM, PTL, PB, KS, AR, VB, XMV, ATW, CS, and SAg contributed to data acquisition, analysis, interpretation, and critically revising the manuscript. SiA, KS, PB, and SAg designed the figures. SiA and SAg drafted the manuscript. AR, CS, and SAg designed the study and acquired funding. All authors have read and approved the current version of the manuscript.

## 7. Conflict of interest statement

The authors declare no competing conflicts of interest regarding this work.

## Notes

### Competing Interest Statement

The authors have declared no competing interest.

