## Supplemental File for "Microfabrication-based engineering of biomimetic dentin-like constructs to simulate dental aging"

**Supplementary figures:**

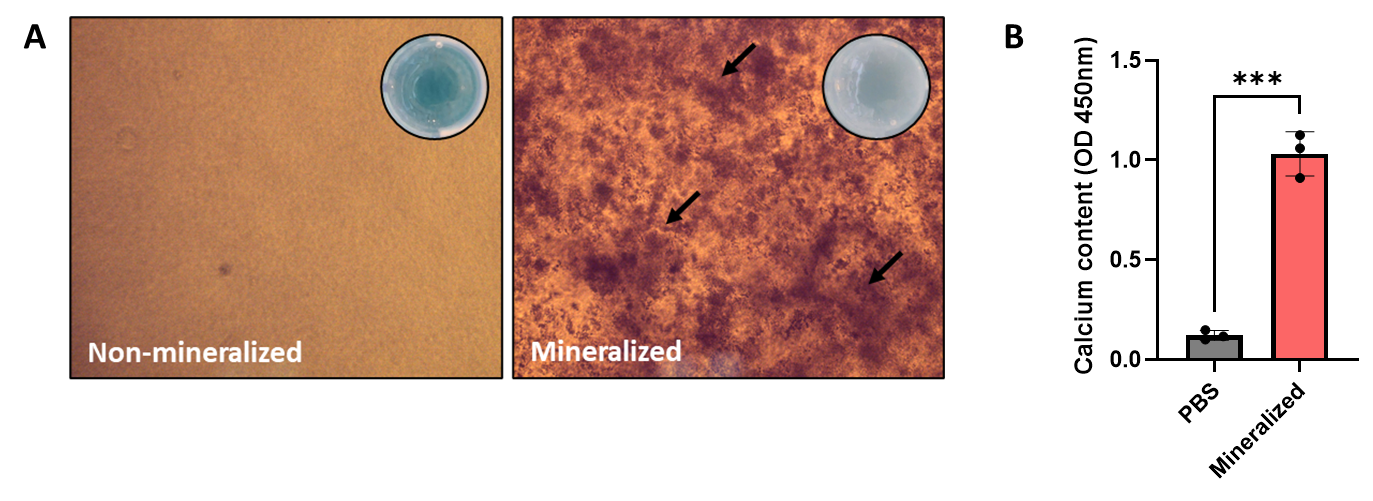
**Microfabrication-based engineering of biomimetic dentin-like constructs to simulate dental aging**

**Supplementary Figure 1: Alizarin Red confirms calcium deposition in type-I collagen substrates.** (A) Phase contrast microscopy of type-I collagen gels displaying calcium deposition (black arrows, 10X magnification). Insets depict collagen in 96-well plates, where the mineralized gels can be observed with white color and reduced translucency. (B) Quantification of calcium content in collagen as a function of Alizarin Red staining (n=3; t-test).

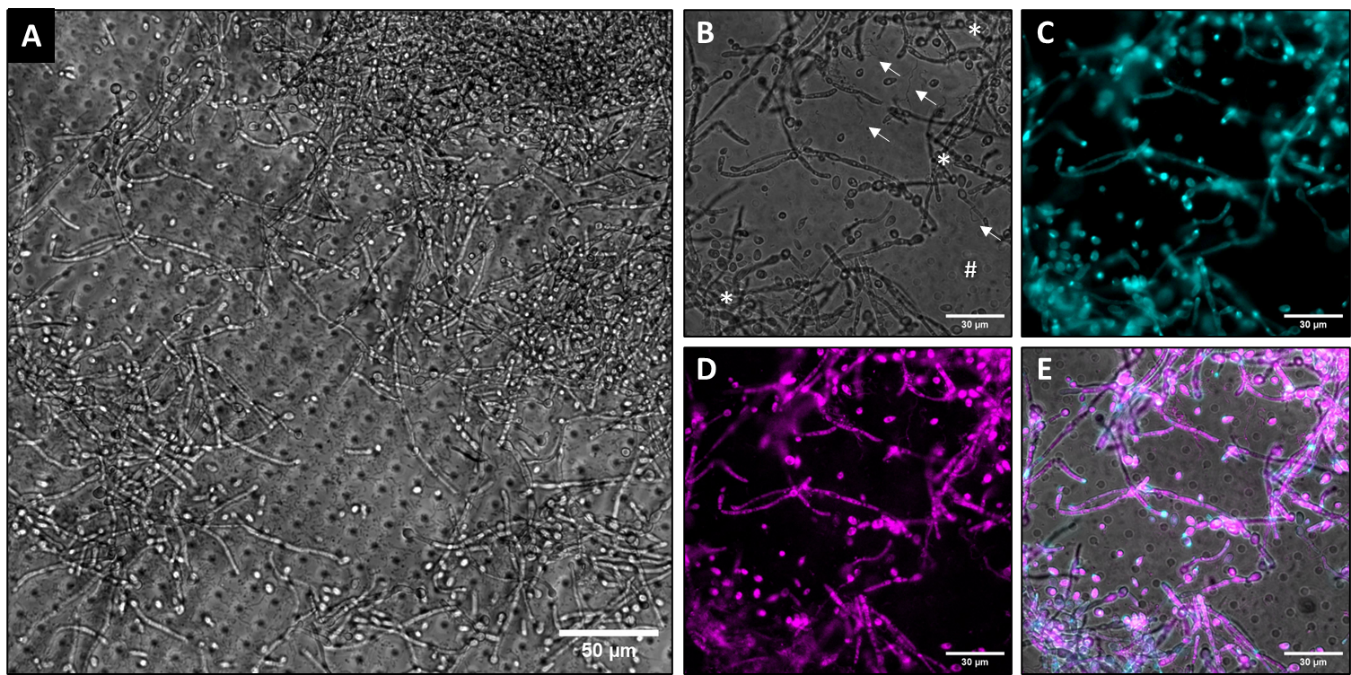

**Supplementary Figure 2: Visualization of dual-species biofilm formation on biomimetic dentin constructs with epifluorescence microscopy.** (A) Phase contrast image of a *C. albicans* and *S. mutans* biofilm on a microfabricated PDMS device. (B) Transmitted light, (C) SYTO9, (D) Calcofluor white, and (E) merged channel images of the dual species biofilm, as a proof-of-concept that the microfabricated substrates can be employed to characterize polymicrobial biofilm formation with an epifluorescence microcopy setup.
